# Predictive computational phenotyping and biomarker discovery using reference-free genome comparisons

**DOI:** 10.1101/045153

**Authors:** Alexandre Drouin, Sébastien Giguère, Maxime Déraspe, Mario Marchand, Michael Tyers, Vivian G. Loo, Anne-Marie Bourgault, François Laviolette, Jacques Corbeil

## Abstract

**Background:** The identification of genomic biomarkers is a key step towards improving diagnostic tests and therapies. We present a reference-free method for this task that relies on a *k*-mer representation of genomes and a machine learning algorithm that produces intelligible models. The method is computationally scalable and well-suited for whole genome sequencing studies.

**Results:** The method was validated by generating models that predict the antibiotic resistance of *C. difficile, M. tuberculosis, P. aeruginosa*, and *S. pneumoniae* for 17 antibiotics. The obtained models are accurate, faithful to the biological pathways targeted by the antibiotics, and they provide insight into the process of resistance acquisition. Moreover, a theoretical analysis of the method revealed tight statistical guarantees on the accuracy of the obtained models, supporting its relevance for genomic biomarker discovery.

**Conclusions:** Our method allows the generation of accurate and interpretable predictive models of phenotypes, which rely on a small set of genomic variations. The method is not limited to predicting antibiotic resistance in bacteria and is applicable to a variety of organisms and phenotypes. Kover, an efficient implementation of our method, is open-source and should guide biological efforts to understand a plethora of phenotypes (http://github.com/aldro61/kover/).

## Background

Despite an era of supercomputing and increasingly precise instrumentation, many biological phenomena remain misunderstood. For example, phenomena such as the development of some cancers, or the lack of efficiency of a treatment on an individual, still puzzle researchers. One approach to understanding such events is the elaboration of *case-control* studies, where a group of individuals that exhibit a given biological state (phenotype) is compared to a group of individuals that do not. In this setting, one seeks biological characteristics (biomarkers), that are predictive of the phenotype. Such biomarkers can serve as the basis for diagnostic tests, or they can guide the development of new therapies and drug treatments by providing insight on the biological processes that underlie a phenotype (Azuaje, 2011; Koboldt et al., 2013; Mbianda et al., 2015; Simon, 2011). With the help of computational tools, such studies can be conducted at a much larger scale and produce more significant results.

In this work, we focus on the identification of genomic biomarkers. These include any genomic variation, from single nucleotide substitutions and indels, to large scale genomic rearrangements. With the increasing throughput and decreasing cost of DNA sequencing, it is now possible to search for such biomarkers in the whole genomes of a large set of individuals (Koboldt et al., 2013; van Dijk et al., 2014). This motivates the need for computational tools that can cope with large amounts of genomic data and identify the subtle variations that are biomarkers of a phenotype.

Genomic biomarker discovery relies on multiple genome comparisons. Genomes are typically compared based on a set of single nucleotide polymorphisms (SNP) (Brookes, 1999; Koboldt et al., 2013; Nielsen et al., 2011). A SNP exists at a single base pair location in the genome when a variation occurs within a population. The identification of SNPs relies on multiple sequence alignment, which is com putationally expensive and can produce inaccurate results in the presence of large-scale genomic rearrangements, such as gene insertions, deletions, duplications, inversions, or translocations (Bonham-Carter et al., 2014; Leimeister et al., 2014; Song et al., 2014; Vinga & Almeida, 2003; Vinga, 2007).

Recently, methods for genome comparison that alleviate the need for multiple sequence alignment, i.e., reference-free genome comparison, have been investigated (Bonham-Carter et al., 2014; Leimeister et al., 2014; Song et al., 2014; Vinga & Almeida, 2003; Vinga, 2007). In this work, we use such an approach, by comparing genomes based on the *k*-mers, i.e., sequences of k nucleotides, that they contain. The main advantage of this method is that it is robust to genomic rearrangements. Moreover, it provides a fully unbiased way of comparing genomic sequences and identifying variations that are associated with a phenotype. However, this genomic representation is far less compact than a set of SNPs and thus poses additional computational challenges.

In this setting, the objective is to find the most concise set of genomic features (*k*-mers) that allows for accurate prediction of the phenotype (Azuaje, 2011). Including uninformative or redundant features in this set would lead to additional validation costs and could mislead researchers. In this work, we favor an approach based on machine learning, where we seek a computational model of the phenotype that is accurate and sparse, i.e. that relies on the fewest genomic features. Learning such models from large data representations, such as the *k*-mer representation, is a challenging problem (Hastie et al., 2013). Indeed, there are many more genomic features than genomes, which increases the danger of overfitting, i.e., learning random noise patterns that lead to poor generalization performance. In addition, the majority of the *k*-mers are uninformative and cannot be used to predict the phenotype. Finally, due to the structured nature of genomes, many *k*-mers occur simultaneously and are thus highly correlated.

Previous work in the field of biomarker discovery has generally combined feature selection and predictive modeling methods (Azuaje, 2011; Saeys et al., 2007). Feature selection serves to identify features that are associated with the phenotype. These features are then used to construct a predictive model with the hope that it can accurately predict the phenotype. The most widespread approach consists in measuring the association between the features and the phenotype with a statistical test, such as the *χ*^2^ test or a t-test. Then, some of the most associated features are selected and given to a modeling algorithm. In the machine learning literature, such methods are referred to as *filter methods* (Guyon & Elisseeff, 2003; Hastie et al., 2013).

When considering millions of features, it is not possible to efficiently perform multivariate statistical tests. Hence, filter methods are limited to univariate statistical tests. While univariate filters are highly scalable, they discard multivariate patterns in the data, that is, combinations of features that are, together, predictive of the phenotype. Moreover, the feature selection is performed independently of the modeling, which can lead to a suboptimal choice of features. *Embedded methods* address these limitations by integrating the feature selection in the learning algorithm (Guyon & Elisseeff, 2003; Saeys et al., 2007). These methods select features based on their ability to compose an accurate predictive model of the phenotype. Moreover, some of these methods, such as the Set Covering Machine (Marchand & Shawe-Taylor, 2002), can consider multivariate interactions between features.

In this study, we propose to apply the Set Covering Machine (SCM) algorithm to genomic biomarker discovery. We devise extensions to this algorithm that make it well suited for learning from extremely large sets of genomic features. We combine this algorithm with the *k*-mer representation of genomes, which reveals uncharacteristically sparse models that explicitly highlight the relationship between genomic variations and the phenotype of interest. We present statistical guarantees on the accuracy of the models obtained using this approach. Moreover, we propose an efficient implementation of the method, which can readily scale to large genomic datasets containing thousands of individuals and hundreds of millions of *k*-mers.

The method was used to model the antibiotic resistance of four common human pathogens, including Gram-negative and Gram-positive bacteria. Antibiotic resistance is a growing public health concern, as many multidrug-resistant bacterial strains are starting to emerge. This compromises our ability to treat common infections, which increases mortality and health care costs (World Health Organization, 2014; Davies & Davies, 2010). Better computational methodologies to assess resistance phenotypes will assist in tracking epidemics, improve diagnosis, enhance treatment, and facilitate the development of new drugs (Earle et al., 2016; Bradley et al., 2015). This study highlights that, with whole genome sequencing and machine learning algorithms, such as the SCM, we can readily zero in on the genes, mutations, and processes responsible for antibiotic resistance and other phenotypes of interest.

## Machine learning for biomarker discovery

The problem of distinguishing two groups of living organisms based on their genomes can be formalized as a *supervised learning* problem. In this setting, we assume that we are given a data sample *𝒮* that contains m *learning examples*. These examples are pairs (x, *y*), where x is a genome and *y* is a label that corresponds to one of two possible phenotypes. More specifically, we assume that x ∈ {*A, T, G,C*}*, which corresponds to the set of all possible strings of DNA nucleotides and that *y* ∈ {0,1}. In this work, the label *y* = 1 is assigned to the case group and *y* = 0 to the control group. The examples in *𝒮* are assumed to be drawn independently from an unknown, but fixed, data generating distribution *D*. Hence 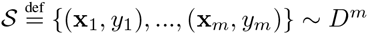.

Most learning algorithms are designed to learn from a vector representation of the data. Thus, to learn from genomes, we must define a function ***φ***: {*A, T, G, C*}* → ℝ^*d*^, that takes a genome as input and maps it to some *d* dimensional vector space (the feature space). We choose to represent each genome by the presence or absence of every possible *k*-mer. This representation is detailed in the Methods section.

Subsequently, a learning algorithm can be applied to the set 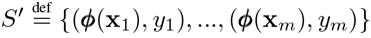 to obtain a model *h*: ℝ^*d*^ → {0,1}. The model is a function that, given the feature representation of a genome, estimates the associated phenotype. The objective is thus to obtain a model *h* that has a good generalization performance, i.e., that minimizes the probability, *R*(*h*), of making a prediction error for any example drawn according to the distribution *D*, where

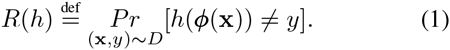

### Application specific constraints

Biomarker discovery leads to two additional constraints on the model *h*. These are justified by the cost of applying the model in practice and on the ease of interpretation of the model by domain experts.

First, we strive for a model that is sparse, i.e., that uses a minimal set of features to predict the phenotype. This property is important, as it can greatly reduce the cost of applying the model in practice. For example, if the model relies on a sufficiently small number of features, these can be measured by using alternative methods, e.g., polymerase chain reaction (PCR), rather than sequencing entire genomes.

In addition, the model must be easily interpretable by domain experts. This is essential for extracting useful biological information from the data, to facilitate comprehension, and is critical for adoption by the scientific community. We make two observations in an attempt to obtain a clear definition of interpretability. The first is that the structure of a model can affect its interpretability. For example, rule-based models, such as decision trees (Breiman et al., 1984), are naturally understood as their predictions consist in answering a series of questions; effectively following a path in the tree. In contrast, linear models, such as those obtained with support vector machines (Cortes & Vapnik, 1995) or neural networks (Cheng & Titterington, 1994), are complex to interpret, as their predictions consist in computing linear combinations of features. The second observation is that, regardless of the structure of the model, sparsity is an essential component in interpretability, since models with many rules are inevitably more tedious to interpret.

### The Set Covering Machine

The SCM (Marchand & Shawe-Taylor, 2002) is a learning algorithm that uses a greedy approach to produce uncharacteristically sparse rule-based models. In this work, the rules are individual units that detect the presence or the absence of a *k*-mer in a genome. These rules are boolean-valued, i.e., they can either output true or false. The models learned by the SCM are logical combinations of such rules, which can be conjunctions (logical-AND) or disjunctions (logical-OR). To predict the phenotype associated with a genome, each rule in the model is evaluated and the results are aggregated to obtain the prediction. A conjunction model assigns the positive class (*y* = 1) to a genome if *all* the rules output true, whereas a disjunction model does the same if *at least one* rule outputs true.

The time required for learning a model with the SCM grows linearly with the number of genomes in the dataset and with the number of *k*-mers under consideration. This algorithm is thus particularly well suited for learning from large genomic datasets. Moreover, as it will be discussed later, we have developed an efficient implementation of the SCM, which can easily scale to hundreds of millions of *k*-mers and thousands of genomes, while requiring a few gigabytes of memory. This is achieved by keeping the data on external storage, e.g., a hard drive, and accessing it in small contiguous blocks. This is in sharp contrast with other learning algorithms, which require that the entire dataset be stored in the computer’s memory.

The SCM algorithm is detailed in Additional File 1 – Appendix 1. In the Methods section, we propose algorithmic and theoretical extensions to the SCM algorithm that make it a method of choice for genomic biomarker discovery.

## Results

### Data

Antibiotic resistance datasets were acquired for four bacterial species: *Clostridium difficile, Mycobacterium tuberculosis*, *Pseudomonas aeruginosa*, and *Streptococcus pneumoniae*. Each dataset was a combination of whole genome sequencing reads and antibiotic susceptibility measurements for multiple isolates. The *M. tuberculosis, P. aeruginosa*, and *S. pneumoniae* data were respectively obtained from Merker et al. (Merker et al., 2015), Kos et al. (Kos et al., 2015) and Croucher et al. (Croucher et al., 2013). The *C. difficile* data were obtained from Dr. Loo and Dr. Bourgault. The genomes were submitted to the European Nucleotide Archive [EMBL:PRJEB11776] and the antibiotic susceptibility measurements are provided in Additional data file 1.

The sequencing data, which are detailed in Additional File 2 - Table S1, were acquired using a variety of Illumina platforms. The genomes were assembled and subsequently split into *k*-mers of length 31 (Methods). Guidelines for selecting an appropriate *k*-mer length are provided in Methods.

Each (pathogen, antibiotic) combination was considered individually, yielding 17 datasets in which the number of examples (*m*) ranged from 111 to 556 and the number of *k*-mers 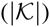 ranged from 10 to 123 millions. Figure 1 shows the distribution of resistant and sensitive isolates in each dataset. The datasets are further detailed in Additional File 2 - Table S2.

**Figure 1.**
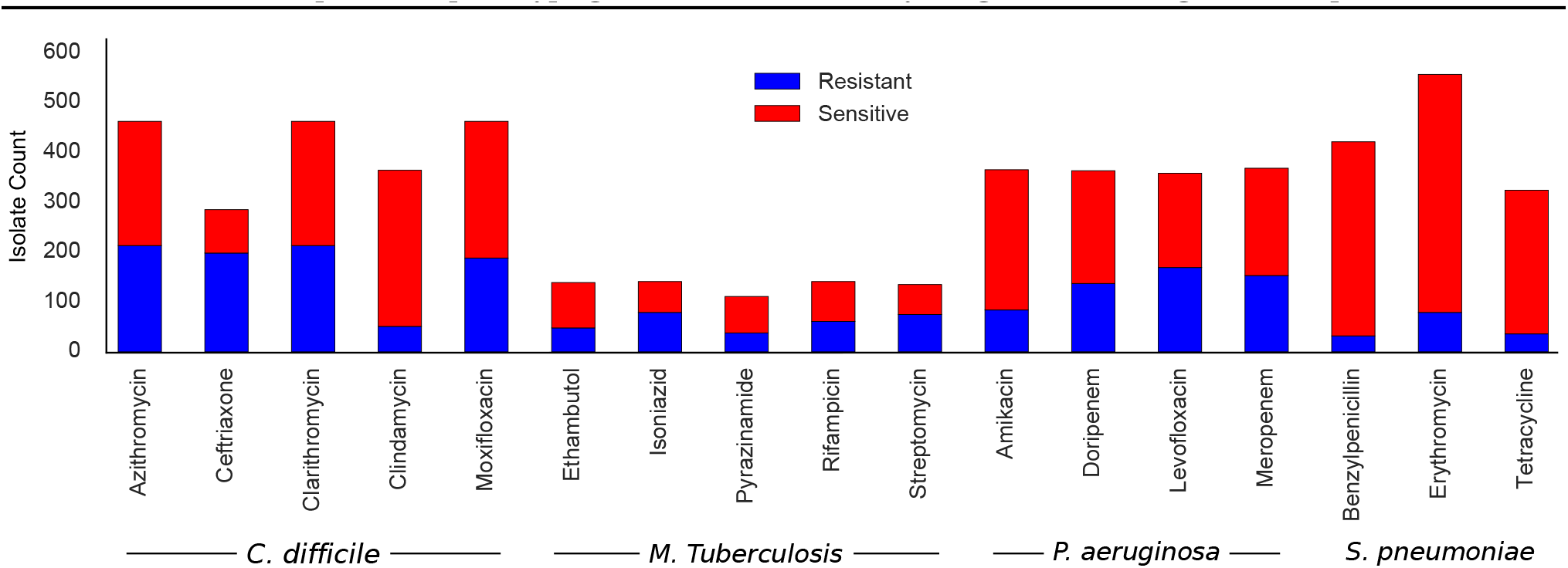
Distribution of resistant and sensitive isolates in each dataset.

### The SCM models are sparse and accurate

The models obtained using the SCM were compared to those obtained using other machine learning algorithms based on their generalization performance and sparsity. Comparisons were made with rule-based models: the CART decision tree algorithm (Breiman et al., 1984), linear classifiers: *L*_1_ -norm and *L*_2_-norm regularized linear support vector machines (L1SVM, L2SVM) (Cortes & Vapnik, 1995), and kernel methods: polynomial and linear kernel support vector machines (PolySVM, LinSVM) (Schölkopf et al., 2004; Shawe-Taylor & Cristianini, 2004). CART and support vector machines are state-of-the-art machine learning algorithms that have been abundantly used in biological applications (Kingsford & Salzberg, 2008; Noble, 2006). Publicly available implementations of these algorithms were used: Scikit-learn (Pedregosa et al., 2011) for CART, LIBLINEAR (Fan et al., 2008) for L1SVM and L2SVM, and LIBSVM (Chang & Lin, 2011) for PolySVM and LinSVM.

The following protocol was used to compare the algorithms. Each dataset was split into a training set (2/3 of the data) and a separate testing set (1/3). 5-fold cross-validation was performed on the training set to select the best hyperparameter values. Finally, each algorithm was trained on the training set and predictions were computed on the held-out testing set. For each algorithm, the generalization performance was measured by the error rate on the independent testing set and sparsity was measured by the number of *k*-mers that contributed to the model. This procedure was repeated 10 times, on different partitions of the data, and the algorithms were compared based on the average error rate and sparsity. A Wilcoxon signed-rank test (Wilcoxon, 1945) was used to assess the statistical significance of the comparisons.

The algorithms were also compared to a baseline method that predicts the most abundant class in the training set (resistant or sensitive).

Our implementation of the SCM was able to learn from the entire feature space, that is, all the *k*-mers. The time required for training the algorithm varied between 33 seconds and two hours, depending on the dataset, and the memory requirements were always inferior to eight gigabytes. In contrast, the CART, L1SVM, and L2SVM algorithms were unable to learn from the entire feature space. For these algorithms, the entire dataset had to be stored in the computer’s memory, generating massive memory requirements. Hence, these algorithms were combined with a feature selection step that reduced the size of the feature space.

**Table 1.** Feature Selection: Average testing set error rate and sparsity (in parentheses) for 10 random partitions of the data. Results are shown for the SCM, which uses the entire feature, and the feature selection-based methods: *χ*^2^ + CART, *χ*^2^ + L1SVM, *χ*^2^ + L2SVM and *χ*^2^ + SCM. The baseline method predicts the most abundant class in the training set. The smallest error rates are in bold.

| Dataset | SCM | $\chi^2$ + CART | $\chi^2$ + L1SVM | $\chi^2$ + L2SVM | $\chi^2$ + SCM | Baseline |
| --- | --- | --- | --- | --- | --- | --- |
| ***C. difficile*** |  |  |  |  |  |  |
| Azithromycin | **0.030** (3.3) | 0.086 (7.2) | 0.064 (20326.0) | 0.056 ($10^6$) | 0.075 (3.0) | 0.446 |
| Ceftriaxone | **0.073** (2.6) | 0.117 (6.8) | 0.087 (8114.1) | 0.102 ($10^6$) | 0.111 (3.2) | 0.306 |
| Clarithromycin | **0.011** (3.0) | 0.070 (8.0) | 0.062 (36686.1) | 0.059 ($10^6$) | 0.069 (3.5) | 0.446 |
| Clindamycin | 0.021 (1.4) | 0.011 (2.0) | 0.009 (598.2) | 0.021 ($10^6$) | **0.008** (2.3) | 0.136 |
| Moxifloxacin | **0.020** (1.0) | **0.020** (1.3) | **0.020** (25.6) | 0.048 ($10^6$) | 0.021 (1.1) | 0.390 |
| ***M. tuberculosis*** |  |  |  |  |  |  |
| Ethambutol | 0.179 (1.4) | 0.185 (1.9) | **0.153** (201.3) | 0.221 ($10^6$) | 0.174 (3.2) | 0.351 |
| Isoniazid | 0.021 (1.0) | 0.021 (1.1) | **0.017** (104.7) | 0.125 ($10^6$) | 0.021 (1.2) | 0.421 |
| Pyrazinamide | **0.318** (3.1) | 0.371 (4.4) | 0.353 (481.2) | 0.342 ($10^6$) | 0.366 (5.8) | 0.347 |
| Rifampicin | 0.031 (1.4) | 0.031 (1.5) | 0.031 (130.0) | 0.196 ($10^6$) | **0.029** (1.3) | 0.452 |
| Streptomycin | 0.050 (1.0) | 0.052 (1.6) | **0.043** (98.8) | 0.137 ($10^6$) | 0.050 (2.1) | 0.435 |
| ***P. aeruginosa*** |  |  |  |  |  |  |
| Amikacin | 0.175 (4.9) | 0.206 (14.1) | 0.187 (11514.6) | **0.164** ($10^6$) | **0.164** (9.7) | 0.216 |
| Doripenem | 0.270 (1.4) | **0.261** (1.9) | **0.261** (950.0) | 0.275 ($10^6$) | 0.307 (8.5) | 0.359 |
| Levofloxacin | **0.072** (1.2) | 0.076 (1.0) | 0.085 (148.9) | 0.212 ($10^6$) | 0.083 (3.5) | 0.463 |
| Meropenem | 0.267 (1.6) | **0.261** (1.0) | 0.328 (5368.5) | 0.327 ($10^6$) | 0.331 (9.1) | 0.404 |
| ***S. pneumoniae*** |  |  |  |  |  |  |
| Benzylpenicillin | 0.013 (1.1) | 0.012 (2.3) | **0.011** (124.9) | 0.013 ($10^6$) | 0.013 (1.3) | 0.073 |
| Erythromycin | **0.037** (2.0) | 0.047 (3.8) | 0.041 (328.8) | 0.042 ($10^6$) | 0.041 (5.1) | 0.142 |
| Tetracycline | 0.031 (1.1) | **0.029** (1.2) | 0.032 (1108.5) | 0.037 ($10^6$) | 0.033 (2.2) | 0.106 |

**Table 2.** Entire feature space: Average testing set error rate and sparsity (in parentheses) for 10 random partitions of the data. Results are shown for the SCM and the kernel methods: LinSVM and PolySVM. The baseline method predicts the most abundant class in the training set. The smallest error rates are in bold.

| Dataset | SCM | LinSVM | PolySVM | Baseline |
| --- | --- | --- | --- | --- |
| ***C. difficile*** |  |  |  |  |
| Azithromycin | **0.030** (3.3) | 0.050 (32 752 570) | 0.048 (32 752 570) | 0.446 |
| Ceftriaxone | **0.073** (2.6) | 0.079 (25 405 987) | 0.076 (25 405 987) | 0.306 |
| Clarithromycin | **0.011** (3.0) | 0.053 (32 752 570) | 0.053 (32 752 570) | 0.446 |
| Clindamycin | **0.021** (1.4) | 0.039 (30 988 214) | 0.039 (30 988 214) | 0.136 |
| Moxifloxacin | **0.020** (1.0) | 0.054 (32 752 570) | 0.048 (32 752 570) | 0.390 |
| ***M. tuberculosis*** |  |  |  |  |
| Ethambutol | **0.179** (1.4) | 0.215 (9 465 489) | 0.221 (9 465 489) | 0.351 |
| Isoniazid | **0.021** (1.0) | 0.117 (9 701 935) | 0.119 (9 701 935) | 0.421 |
| Pyrazinamide | **0.318** (3.1) | 0.382 (8 058 479) | 0.382 (8 058 479) | 0.347 |
| Rifampicin | **0.031** (1.4) | 0.200 (9 701 935) | 0.204 (9 701 935) | 0.452 |
| Streptomycin | **0.050** (1.0) | 0.143 (9 282 080) | 0.148 (9 282 080) | 0.435 |
| ***P. aeruginosa*** |  |  |  |  |
| Amikacin | **0.175** (4.9) | 0.184 (116 441 834) | 0.179 (116 441 834) | 0.216 |
| Doripenem | **0.270** (1.4) | 0.288 (122 438 059) | 0.281 (122 438 059) | 0.359 |
| Levofloxacin | **0.072** (1.2) | 0.221 (122 216 859) | 0.225 (122 216 859) | 0.463 |
| Meropenem | **0.267** (1.6) | 0.329 (123 466 989) | 0.331 (123 466 989) | 0.404 |
| ***S. pneumoniae*** |  |  |  |  |
| Benzylpenicillin | **0.013** (1.1) | 0.015 (8 968 176) | 0.015 (8 968 176) | 0.073 |
| Erythromycin | **0.037** (2.0) | 0.046 (9 666 898) | 0.047 (9 666 898) | 0.142 |
| Tetracycline | **0.031** (1.1) | 0.039 (8 657 259) | 0.037 (8 657 259) | 0.106 |

The next two sections compare the SCM to two categories of methods: those that require feature selection and those that learn from the entire feature space. In both cases, the SCM was found to yield the sparsest and most accurate models.

#### Feature selection

Feature selection was performed using a univariate filter that measured the association between each feature and the phenotype (Azuaje, 2011; Guyon & Elisseeff, 2003; Saeys et al., 2007). Using the *χ*^2^ test of independence, the 1000 000 most associated features were retained. Results comparing the SCM, which uses all features, to the univariately filtered algorithms: *χ*^2^ + CART, *χ*^2^ + L1SVM, and *χ*^2^ + L2SVM, are shown in Table 1.

In terms of error rate, all the algorithms surpass the baseline method, indicating that relevant information about antibiotic resistance was found in the genomes. The error rate of the SCM is smaller or equal to that of *χ*^2^ + CART on 12/17 dataset (*p* = 0.074), *χ*^2^ + L1SVM on 11/17 datasets (*p* = 0. 179), and *χ*^2^ + L2SVM on 16/17 datasets (*p* = 0.001). Moreover, the SCM was found to learn sparser models than these algorithms (*χ*^2^ + CART: *p* = 0.003, *χ*^2^ + L1SVM: *p* = 0.0003, *χ*^2^ + L2SVM: *p* = 0.0003).

In addition, the SCM was compared to a variant which uses univariate feature selection (*χ*^2^ + SCM). This comparison revealed that the SCM surpasses the *χ*^2^ + SCM in terms of accuracy (*p* = 0.001) and sparsity (*p* = 0.054), highlighting the importance of multivariate patterns in the data (Table 1).

The ability to consider the entire feature space is thus critical and eliminates the selection biases of feature selection methods. However, for most machine learning algorithms, this remains impossible due to computational limitations and the danger of overfitting. The next section compares the SCM to two methods that learn from the entire feature space.

#### Entire feature space

By means of the kernel trick, kernel methods can efficiently learn from very high dimensional feature spaces (Schölkopf et al., 2004; Shawe-Taylor & Cristianini, 2004). However, as opposed to the SCM, they do not yield sparse models that can be interpreted by domain experts.

The SCM was compared to support vector machines coupled with linear (LinSVM) and polynomial (PolySVM) kernels. When learning from our binary genomic representation (Methods), the LinSVM yields a linear model that considers the presence or absence of each *k*-mer. Moreover, the PolySVM yields a linear model that considers all possible conjunctions of 1 to d *k*-mers, where d is a hyperparameter of the kernel. The obtained models are thus akin to those of the SCM, making this comparison particularly interesting.

The results, shown in Table 2, indicate that the SCM models are both more accurate (LinSVM: *p* = 0.0003, PolySVM: *p* = 0.0003) and sparser (LinSVM: *p* = 0.001, PolySVM: *p* = 0.006) than those of the aforementioned algorithms. Further analysis revealed that the poor performance of LinSVM and PolySVM is due to overfitting (Additional File 2 - Table S3), which likely occurs due to the immensity of the feature space. In contrast, the SCM was not found to overfit. This is consistent with the theoretical result described in Methods, which indicates that the SCM is not prone to overfitting in settings where the number of features is much larger than the number of examples.

In summary, it is not only its ability to consider the entire feature space, but also its sparsity and high resistance to overfitting that make for the strong performance of the SCM. In complement to these results, the mean and standard deviation of the sensitivity, specificity, and error rate for each algorithm are provided in Additional File 2 - Tables S4, S5, S6.

### The SCM models are biologically relevant

The biological relevance of the SCM models was investigated. To achieve this, the algorithm was retrained on each dataset, using all the available data. This yielded a single phenotypic model for each dataset. Then, the *k*-mer sequences of the rules in the models were annotated by using Nucleotide BLAST (Altschul et al., 1990) to search them against a set of annotated genomes.

Moreover, for each rule in the models, rules that the SCM found to be equally predictive of the phenotype (equivalent rules) were considered in the analysis. Such rules are not used for prediction, but can provide insight on the type of genomic variation that was identified by the algorithm (see Methods). For example, a small number of rules targeting *k*-mers that all overlap on a single or few nucleotides, suggests a point mutation. Alternatively, a large number of rules, that target *k*-mers which can be assembled to form a long sequence, suggests a large-scale genomic variation, such as a gene insertion or deletion.

The annotated models for each dataset are illustrated in Additional File 3 - Figure S1. Below, a subset of these models, which is illustrated in Figure 2, is discussed. For each genomic variation identified by the algorithm, a thorough literature review was performed, with the objective of finding known, and validated, associations to antibiotic resistance.

**Figure 2.**
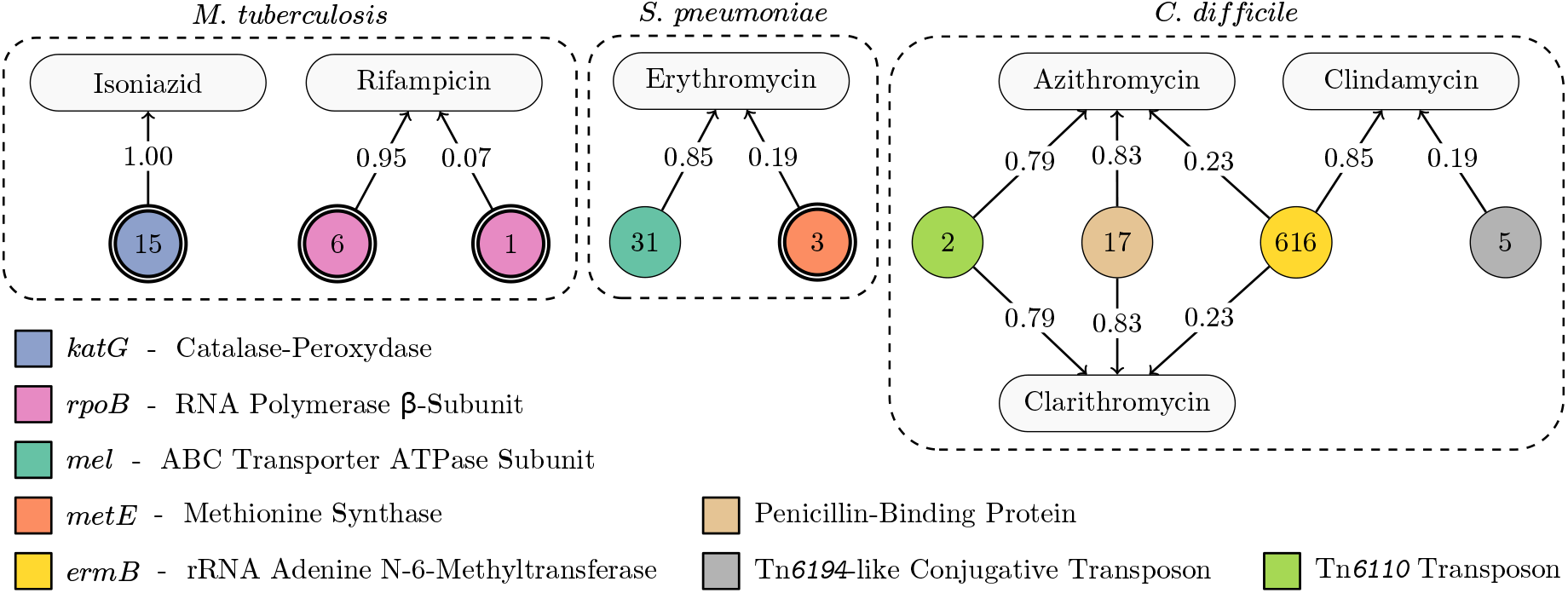
Six antibiotic resistance models, which are all disjunctions (logical-OR). The rounded rectangles correspond to antibiotics. The circular nodes correspond to *k*-mer rules. A single border indicates a presence rule and a double border indicates an absence rule. The numbers in the circles show to the number of equivalent rules. A rule is connected to an antibiotic if it was included in its model. The weight of the edges gives the importance of each rule as defined by Equations (3) and (4). The models for all 17 datasets are illustrated in Additional File 3 - Figure S1.

For *M. tuberculosis*, the isoniazid resistance model contains a single rule which targets the *katG* gene. This gene encodes the catalase-peroxidase enzyme (KatG), which is responsible for activating isoniazid, a prodrug, into its toxic form. As illustrated in Figure 3, the *k*-mers associated with this rule and its equivalent rules all overlap a concise locus of *katG*, suggesting the occurrence of a point mutation. This locus contains codon 315 of KatG, where mutations S315G, S315I, S315N and S315T are all known to result in resistance (Cade et al., 2010; Da Silva & Palomino, 2011). A multiple sequence alignment revealed that these variants were all present in the dataset. The SCM therefore selected a rule that captures the absence of the wild-type sequence at this locus, effectively including the presence of all the observed variants.

**Figure 3.**
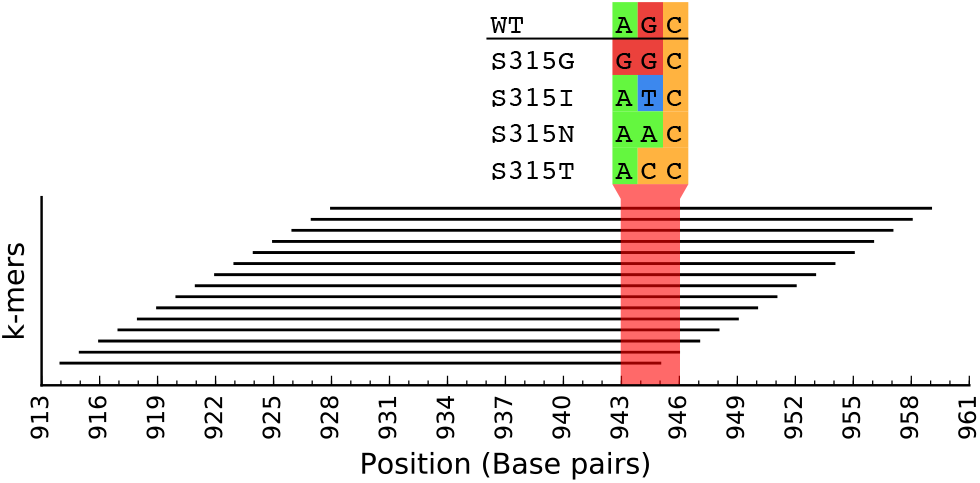
Going beyond *k*-mers:

This figure shows the location, on the *katG* gene, of each *k*-mer targeted by the isoniazid model (rule and equivalent rules). All the *k*-mers overlap a concise locus, suggesting that it contains a point mutation that is associated with the phenotype. A multiple sequence alignment revealed a high level of polymorphism at codon 315 (shown in red). The wildtype sequence (WT), as well as the resistance conferring variants S315G, S315I, S315N and S315T, were observed. The rule in the model captures the absence of WT and thus, includes the occurrence of all the observed variants.

The rifampicin resistance model contains two rules, which target the rifampicin resistance-determining region (RRDR) of the *rpoB* gene. This gene, which encodes the *β*-subunit of the RNA polymerase, is the target of rifampicin. The antibiotic binds to RpoB, which inhibits the elongation of messenger RNA. Mutations in the RRDR are known to cause conformational changes that result in poor binding of the drug and cause resistance (Da Silva & Palomino, 2011). Furthermore, one of the rules has a much greater importance than the other. This suggests the existence of two clusters of rifampicin resistant strains, one being predominant, while both harbor mutations in different regions of the RRDR.

For *S. pneumoniae*, the first and most important rule of the erythromycin resistance model targets the *mel* gene. The *mel* gene is part of the macrolide efflux genetic assembly (MEGA) and is known to confer resistance to erythromycin (Daly et al., 2004; Ambrose et al., 2005). Of note, this gene is found on an operon with either the *mefA* or the *mefE* gene, which are also part of the MEGA and associated with erythromycin resistance (Daly et al., 2004). It is likely that the algorithm targeted the *mel* gene to obtain a concise model that includes all of these resistance determinants. The second rule in the model is an absence rule that targets the wild-type version of the *metE* gene. This gene is involved in the synthesis of methionine (Basavanna et al., 2013). Alterations in this gene could lead to a lack of methionine in the cell and impact the ribosomal machinery, which is the drug’s target. However, further validation is required to confirm this resistance determinant.

For *C. difficile*, the resistance models for azithromycin and clarithromycin, two macrolide antibiotics, share a rule with the resistance model for clindamycin, a lincosamide antibiotic. These three antibiotics function by binding the 50S subunit of the ribosome and interfering with bacterial protein synthesis (Tenson et al., 2003). Cross-resistance between macrolide and lincosamide antibiotics is caused by the presence of the *ermB* gene that encodes rRNA adenine N-6-methyltransferase, an enzyme that methylates position 2058 of the 23S rRNA within the larger 50S subunit (Farrow et al., 2000; Tenson et al., 2003; Vester & Douthwaite, 2001). The shared rule for the macrolide and the lincosamide models rightly targets the *ermB* gene. This rule has 616 equivalent rules, all of the *presence* type, targeting *ermB*. Arguably, the algorithm correctly found the presence of this gene to be a cross-resistance determinant, in agreement with the literature (Farrow et al., 2000; Tenson et al., 2003; Vester & Douthwaite, 2001).

Azithromycin and clarithromycin have similar mechanisms of action and, as expected, their resistance models are identical. They contain a presence rule that targets a region of the Tn*6110* transposon, characterized in *C. difficile* strain QCD-6626 (Brouwer et al., 2011). This region is located 136 base pairs downstream of a 23S rRNA methyl-transferase, which is a gene known to be associated with macrolide resistance (Kaminska et al., 2010). The next rule in the models targets the presence of the penicillin-binding protein, which plays a role in resistance to *β*-lactam antibiotics, such as ceftriaxone (Waxman & Strominger, 1983). Among the azithromycin-resistant isolates in our dataset, 92.7% are also resistant to ceftriaxone. Similarly, 92.2% of the clarithromycin-resistant isolates are resistant to ceftriaxone. Hence, this rule was likely selected due to these strong correlations.

Finally, clindamycin resistance model contains a rule targeting a Tn*6194*-like conjugative transposon. This transposon contains the *ermB* gene, which is associated with resistance to this antibiotic (Wasels et al., 2013). Moreover, it is rarely found in clinical isolates, which could explain its smaller importance.

### Spurious correlations can be overcome

One limitation of statistical methods that derive models from data is their inability to distinguish causal variables from those that are highly correlated with them. To our knowledge, it is very difficult to prevent this pitfall. However the interpretability and the sparsity of the obtained models can be leveraged to identify and circumvent spurious correlations.

One notable example of such a situation is the strong correlation in resistance to antibiotics that do not share common mechanisms of action. These correlations might originate from treatment regimens. For instance, Figure 4 A shows, for *M. tuberculosis*, the proportion of isolates that are identically labeled (resistant or sensitive) for each pair of antibiotics. More formally, this figure shows a matrix *C*, where each entry *C_ij_* corresponds to a pair of datasets 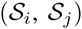 and

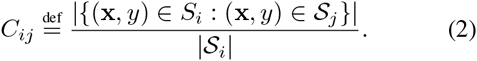

Notice the large proportion of isolates in the streptomycin dataset that are identically labeled in the isoniazid dataset (95.6%). Consequently, the models obtained for streptomycin and isoniazid resistance are identical (Additional File 3 - Figure S1). However, these antibiotics have different mechanisms of action and thus, different resistance mechanisms.

**Figure 4.**
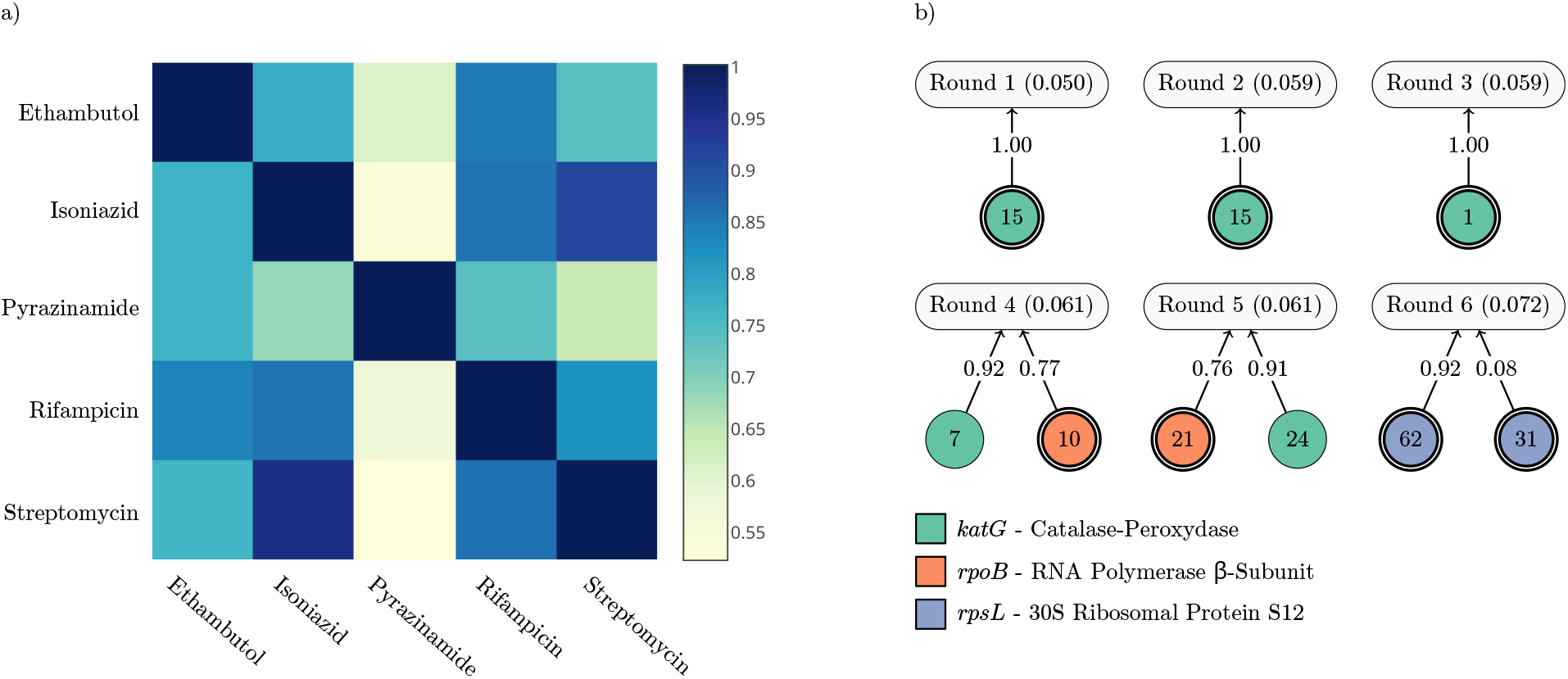
Overcoming spurious correlations: This figures shows how spurious correlations in the *M. tuberculosis* data affect the models produced by the Set Covering Machine. a) For each *M. tuberculosis* dataset, the proportion of isolates that are identically labeled in each other dataset is shown. This proportion is calculated using Equation (2). b) The antibiotic resistance models learned by the SCM at each iteration of the correlation removal procedure. Each model is represented by a rounded rectangle identified by the round number and the estimated error rate. All the models are disjunctions (logical-OR). The circular nodes correspond to *k*-mer rules. A single border indicates a presence rule and a double border indicates an absence rule. The numbers in the circles show to the number of equivalent rules. A rule is connected to an antibiotic if it was included in its model. The weight of the edges gives the importance of each rule.

The following procedure is proposed to eliminate spurious correlations and identify causal genomic variants:

1. Learn a model using the SCM.
2. Validate the association between the rules in the model and the phenotype using mutagenesis and phenotypic assays.
3. If a rule is not rightly associated with the phenotype, remove the *k*-mers of the rule and its equivalent rules from the data.
4. Repeat until a causal association is found.

In practice, the models can be validated by genetically engineering mutants that match the *k*-mer variations targeted by the model. Such mutants can be engineered by diverse means, such as homologous recombination, the CRISPR-Cas9 approach (Hsu et al., 2014), or standard molecular biology cloning. For a conjunction, a multilocus mutant can be engineered to test the synergy between the presence/absence of the *k*-mers. For a disjunction, the rules must be validated individually, by engineering one mutant for each rule in the model. Finally, the phenotypes of the mutants can be experimentally validated using phenotypic assays. For example, antibiotic resistance can be validated by using standard susceptibility testing protocols in the presence of the antibiotic.

Figure 4 B shows a proof of concept, where the iterative procedure was applied to streptomycin resistance. Resistance to this antibiotic is well documented and thus, a literature review was used in lieu of the experimental validation of mutants. Six rounds were required in order to converge to a known resistance mechanism, i.e., the *rpsL* gene (Nair et al., 1993). The models obtained throughout the iterations contained rules targeting the *katG* and the *rpoB* genes, which are respectively isoniazid and rifampicin resistance determinants (Cade et al., 2010; Da Silva & Palomino, 2011). Again, this occurs due to the large proportion of isolates in the streptomycin dataset that are identically labeled in the isoniazid (95.6%) and rifampicin datasets (85.9%).

Hence, should the algorithm identify variations that are correlated with, but not causal of the phenotype, one could detect and eliminate them, eventually converging to causal variants. The search for causality is therefore a feedback between machine learning and experimental biology, which is made possible by the high sparsity and interpretability of the models generated using the SCM.

### The SCM can predict the level of resistance

To further demonstrate how the SCM can be used to explore the relationship between genotypes and phenotypes, it was used to predict the level of benzylpenicillin resistance in *S. pneumoniae*. For this bacterium, penicillin resistance is often mediated by alterations that reduce the affinity of penicillin-binding proteins (Fani et al., 2014). Moderate-level resistance is due to alterations in PBP2b and PBP2x, whereas high-level resistance is due to additional alterations in PBP1a. Based on the antibiotic susceptibility data described in Additional File 2 - Table S2, three levels of antibiotic resistance were defined and used to group the isolates: high-level resistance (R), moderate-level resistance (I) and sensitive (S). We then attempted to discriminate highly resistant isolates from sensitive isolates and moderately resistant isolates from sensitive isolates. The same protocol as in the previous sections was used.

An error rate of 1.3% was obtained for discriminating highly resistant and sensitive isolates. The obtained model correctly targeted the *pbp1a* gene. Based on the protocol presented in Additional File 1 – Appendix 2, all the *k*-mers located in this gene were removed and the experiment was repeated. This yielded a model with an error rate of 1.7% that targeted the *pbp2b* gene. These results are consistent with the literature, since they indicate that alterations in both genes are equally predictive of a high-level of resistance and thus, that they occur simultaneously in isolates that are highly resistant to penicillin (Fani et al., 2014).

An error rate of 6.4% was obtained for discriminating moderately resistant and sensitive isolates. The obtained model correctly targeted the *pbp2b* gene. Again, all the *k*-mers located in this gene were removed and the experiment was repeated. The obtained model had an error rate of 7.2% and targeted the *pbp2x* gene. In accordance with the literature, this indicates that alterations in both genes are predictive of moderate-level resistance. However, our results indicate that alterations in *pbp2b* are slightly more predictive of this phenotype.

## Discussion

We have addressed the problem of learning computational phenotyping models from whole genome sequences. We sought a method that produces accurate models that are interpretable by domain experts, while relying on a minimal set of biomarkers. Our results for predicting antibiotic resistance demonstrate that this goal has been achieved.

Biologically relevant insight was acquired for drug resistance phenotypes. Indeed, within hours of computation, we have retrieved antibiotic resistance mechanisms that have been reported over the past decades. Of note, we have shown that the *k*-mers in the SCM models can be further refined to determine the type of the underlying genomic variations. Hence, this method could be used to rapidly gain insight on the causes of resistance to new antibiotics, for which the mechanism of action might not be fully understood. Furthermore, as our results suggest, our method could be used to discover resistance mechanisms that are shared by multiple antibiotics, which would allow the development of more effective combination therapies.

In terms of accuracy, the method was shown to outperform a variety of machine learning-based biomarker discovery methods. For a majority of datasets, the achieved error rates are well below 10%. Given the inherent noise in antibiotic susceptibility measurements, it is likely that these error rates are near optimal. For *M. tuberculosis* and *P. aeruginosa*, some datasets were shown to have contrasting results, where none of the evaluated methods produced accurate models. A FastQC (and, 2010) analysis revealed that, of the four species considered, these two species have the lowest sequencing data quality (Additional File 2 - Table S1). Moreover, for these species only, the data were acquired using a combination of MiSeq and HiSeq instruments, which could undermine the comparability of the genomes (Loman et al., 2012).

We therefore hypothesize that the inability to learn accurate models on some datasets is either due to the quality of the sequencing data, an insufficient number of learning examples, or extra-genomic factors that influence the phenotype. For instance, epigenetic modifications have been shown to alter gene expression in bacteria and play a role in virulence (Adam et al., 2008; Casadesús & Low, 2006). Assuming the availability of the data, future work could explore extensions to jointly learn models from genetic and epigenetic data.

In terms of sparsity, the SCM was shown to produce the sparsest models. Notably, this was achieved without negatively impacting the prediction accuracy of the models. We hypothesize that this is due to the small number of genomic variations that drive some genome-related phenotypes.

Hence, we presented empirical evidence that, in the context of genomic biomarker discovery, the SCM outperforms a variety of machine learning algorithms, which were selected to have diverse model structures and levels of sparsity. This suggests that the conjunctions and disjunctions produced by the SCM, in addition to being intuitively understandable, are more suitable for this task. In Methods, we provide tight statistical guarantees on the accuracy of the models obtained using our approach. Such theoretical results are uncommon for this type of tool and, together with the empirical results, indicate the SCM is a tool of choice for genomic biomarker discovery.

## Conclusions

The identification of genomic biomarkers is a key step towards improving diagnostic tests and therapies. In this study, we have demonstrated how machine learning can be used to identify such biomarkers in the context of case-control studies. We proposed a method that relies on the Set Covering Machine algorithm to generate models that are accurate, concise and intelligible to domain experts. The obtained models make phenotypic predictions based on the presence or absence of short genomic sequences, which makes them well-suited for translation to the clinical settings using methods such as PCR. The proposed method is broadly applicable and is not limited to predicting drug response in bacteria. Hence, we are confident that this work will transpose to other organisms, phenotypes, and even to scenarios involving complex mixtures of genomes, such as metagenomic studies. The efficiency and the simplicity of the models obtained using our method could guide biological efforts for understanding a plethora of phenotypes.

To facilitate the integration of our method in genomic analysis pipelines, we provide Kover, an implementation that efficiently combines the modeling power of the Set Covering Machine with the versatility of the *k*-mer representation. The implementation is open-source and is available at http://github.com/aldro61/kover.

## Methods

### Genome assembly and fragmentation into *k*-mers

All genomes were assembled using the SPAdes genome assembler (Bankevich et al., 2012) and were subsequently split into *k*-mers using the Ray Surveyor tool, which is part of the Ray *de novo* genome assembler (Boisvert et al., 2010; 2012). Genome assembly is not mandatory for applying our method. Instead, one could use *k*-mer counting software to identify the *k*-mers present in the raw reads of each genome. However, with sufficient coverage, genome assembly can increase the quality of the *k*-mer representation by eliminating sequencing errors. This reduces the number of unique *k*-mers and thus, the size of the feature space.

### Choosing the *k*-mer length

The *k*-mer length is an important parameter of the proposed method. Exceedingly small values of *k* will yield *k*-mers that ambiguously map to multiple genomic loci. Yet, an exceedingly large *k* will yield very specific *k*-mers that only occur in few genomes. To our knowledge, a general protocol for selecting the *k*-mer length does not exist. We therefore propose two approaches to selecting an appropriate length.

The first consists of using prior biological knowledge about the organism under study. For instance, the mutation rate is an important factor to consider. If it is expected to be high (e.g., viruses), small *k*-mers are preferable. Conversely, if the mutation rate is low, longer *k*-mers can be used, allowing the identification of additional genomic variations, such as DNA tandem repeats, which can be relevant for predicting the phenotype (Zhou et al., 2014). Extensive testing has shown that *k* = 31 appears to be optimal for bacterial genome assembly (Boisvert et al., 2012) and recent studies have employed it for reference-free bacterial genome comparisons (Earle et al., 2016; Bradley et al., 2015). Hence, this value was used in the current study.

The second method is better suited for contexts where no prior knowledge is available. It consists of considering *k* as a hyperparameter of the learning algorithm and setting its value by cross-validation. In this case, the algorithm is trained using various values of *k* and the best value is selected based on the cross-validation score. This process is more computationally intensive, since the algorithm needs to be trained multiple times. However, it ensures that the *k*-mer length is selected based on the evidence of a good generalization performance.

In this study, both approaches were compared and shown to yield similar results. Indeed, no significant variation in accuracy was observed for the models obtained with *k* = 31 and with *k* selected from {15,21,31,51,71,91} by cross-validation (Additional File 1 – Appendix 3). This corroborates that *k*-mers of length 31 are well-suited for bacterial genome comparisons. Moreover, it indicates that cross-validation can recover a good *k*-mer length in the absence of prior knowledge.

### Applying the Set Covering Machine to genomes

We represent each genome by the presence or absence of each possible *k*-mer. There are 4^*k*^ possible *k*-mers and hence, for *k* = 31, we consider 4^31^ > 4 · 10^18^ *k*-mers. Let *K* be the set of all, possibly overlapping, *k*-mers present in at least one genome of the training set 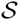. Observe that 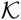 omits *k*-mers that are absent in 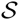 and thus non-discmninatory, which allows the SCM to efficiently work in this enormous feature space. Then, for each genome x, let 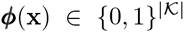 be a 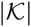 dimensional vector, such that its component *φ_i_*(*x*) = 1 if the *i*-th *k*-mer of 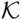 is present in x and 0 otherwise. An example of this representation is given in Figure 5. We consider two types of boolean-valued rules: presence rules and absence rules, which rely on the vectors *φ*(*x*) to determine their outcome. For each *k*-mer 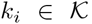, we define a presence rule as 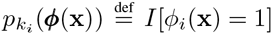 and an absence rule as 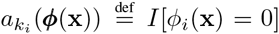, where *I*[*a*] = 1 if *a* is true and *I*[*a*] = 0 otherwise. The SCM, which is detailed in Additional File 1 – Appendix 1, can then be applied by using 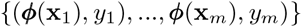 as the set 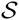 of learning examples and by using the set of presence/absence rules defined above as the set 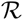 of boolean-valued rules. This yields a phenotypic model which explicitly highlights the importance of a small set of *k*-mers. In addition, this model has a form which is simple to interpret, since its predictions are the result of a simple logical operation.

**Figure 5.**
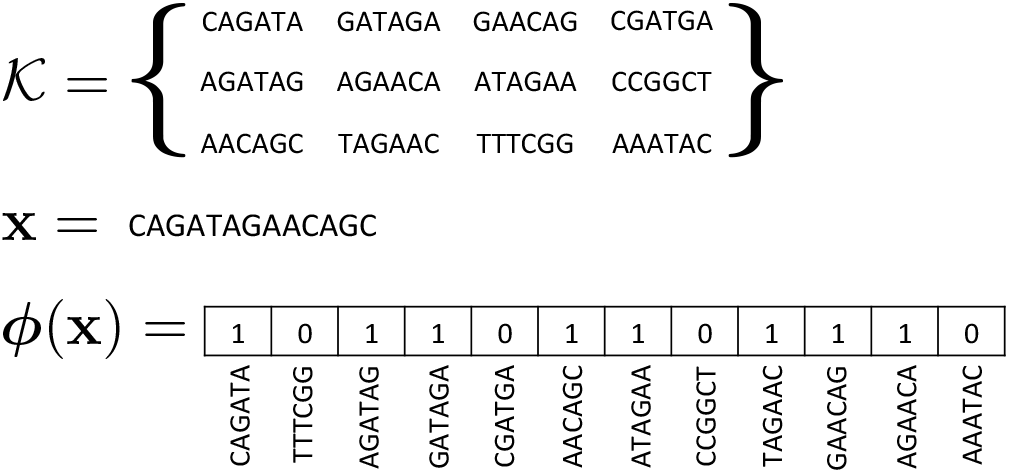
The *k*-mer representation: An example of the *k*-mer representation. Given the set of observed *k*-mers 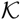 and a genome x. the corresponding vector representation is given by ***φ***(x).

### Tiebreaker function

At each iteration of the SCM algorithm (Marchand & Shawe-Taylor, 2002), the rules are assigned a utility score based on their ability to classify the examples for which the outcome of the model is not settled. The number of such examples decreases at each iteration. Consequently, it is increasingly likely that many rules have an equal utility score. This phenomenon is accentuated when considering many more rules than learning examples, which is the case of biomarker discovery. We therefore extend the algorithm by introducing a tiebreaker function for rules of equal utility. The tiebreaker consists in selecting the rule that best classifies all the learning examples, i.e., the one with the smallest empirical error rate. This simple strategy favors rules that are more likely to be associated with the phenotype.

### Exploiting equivalent rules

When applied to genomic data, the tiebreaker does not always identify a single best rule. This is a consequence of the inherent correlation that exists between *k*-mers that occur simultaneously in the genome, such as *k*-mers that overlap or that are nearby in the genomic structure. The rules that the tiebreaker cannot distinguish are deemed equivalent. Our goal being to obtain concise models, only one of these rules is included in the model and used for prediction. This rule is selected randomly, but other strategies could be applied. As it has been demonstrated in the results, these rules provide a unique approach for deciphering, *de novo*, new biological mechanisms without the need for prior information. Indeed, the set of *k*-mers targeted by these rules can be analyzed to draw conclusions on the type of genomic variation that was identified by the algorithm, e.g., point mutation, indel or structural variation.

### Measuring the importance of rules

We propose a measure of importance for the rules in a conjunction or disjunction model. Taking rule importance into consideration can facilitate the interpretation of the model. Importance should be measured proportionally to the impact of each rule on the predictions of the model. Observe that for any example x, a conjunction model predicts *h*(x) =0 if at least one of its rules returns 0. Thus, when a rule returns 0, it directly contributes to the outcome of the model. Moreover, a conjunction model predicts *h*(x) = 1 if and only if exactly all of its rules return 1. Hence, in this case, all the rules contribute equally to the prediction and thus, we do not need to consider this case in the measure of importance. The importance of a rule r in a conjunction model is therefore given by:

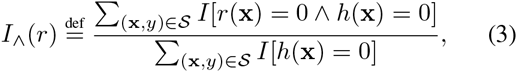

where *r*(x) is the outcome of rule *r* on example x. In contrast, a disjunction model predicts *h*(x) = 1 if at least one of its rules return 1. Moreover, it predicts *h*(x) =0 if and only if exactly all of its rules returns 0. The importance of a rule in a disjunction model is thus given by:

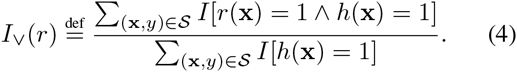

### An upper bound on the error rate

When the number of learning examples is much smaller than the number of features, many machine learning algorithms tend to overfit the training data and thus, have a poor generalization performance (Hastie et al., 2013). Genomic biomarker discovery fits precisely in this regime. Using sample-compression theory (Floyd & Warmuth, 1995; Littlestone & Warmuth, 1986; Marchand & Sokolova, 2005), we obtained an upper bound on the error rate, *R*(*h*), of any model, *h*, learned using our proposed approach. Interestingly, this bound indicates that we are not in a setting where the SCM is prone to overfitting, even if the number of features is much larger than the number of example.

Formally, for any distribution *D*, with probability at least 1 – δ, over all datasets 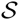 drawn according to *D^m^*, we have that all models *h* have *R*(*h*) ≤ ϵ, where

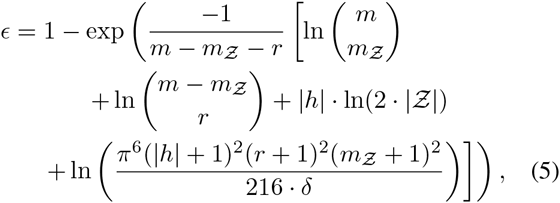

where *m* is the number of learning examples, |*h*| is the number of rules in the model, 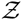 is a set containing 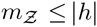 learning examples (genomes) in which each *k*-mer in the model can be found, 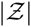 is the total number of nucleotides in 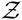 and *r* is the number of prediction errors made by *h* on 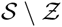. The steps required to obtain this bound are detailed in Additional File 1 – Appendix 4.1.

This theoretical result guarantees that our method will achieve good generalization, regardless of the number of possible features under consideration (4^*k*^), provided that we obtain a sparse model (small |*h*|) that makes few errors on the training set (small *r*). Hence, the occurrence of overfitting is not influenced by the immensity of the feature space under consideration. This is theoretical evidence that the SCM is a method of choice for genomic biomarker discovery studies. Moreover, this is reflected in our empirical results, which indicate that using various *k*-mer lengths, and thus feature spaces of various sizes, does not significantly affect the accuracy of the obtained models (*p* = 0.551) (Additional File 1 – Appendix 3).

This result is counter-intuitive with respect to classical machine learning theory and highlights the benefits of using sample-compression theory to analyze the behavior of learning algorithms.

Finally, following the idea of Marchand and Shawe-Taylor (Marchand & Shawe-Taylor, 2002), we attempted to use the bound value as a substitute for 5-fold crossvalidation. In this case, the bound value was used to determine the best combination of hyperparameter values (Additional File 1 – Appendix 4.2). This led to a sixfold decrease in the number of times the SCM had to be trained and yielded sparser models (*p* = 0.014) with similar accuracies (*p* = 0.463).

### Efficient implementation

The large size of genomic datasets tends to surpass the memory resources of modern computers. Hence, there is a need for algorithms that can process such datasets without solely relying on the computer’s memory. *Out-of-core* algorithms achieve this by making efficient use of external storage, such as file systems. Along with this work, we propose *Kover*, an out-of-core implementation of the Set Covering Machine tailored for presence/absence rules of *k*-mers. Kover implements all the algorithmic extensions proposed in this work. It makes use of the HDF5 library (The HDF Group, 2015) to efficiently store the data and process it in blocks. Moreover, it exploits atomic CPU instructions to accelerate computations. The details are provided in Additional File 1 – Appendix 5. Kover is implemented in the Python and C programming languages, is open-source software and is available free of charge.

### Future work

The proposed method is currently limited to the presence or absence of *k*-mers. This binary representation leads to desirable algorithmic properties and allows the use of highly efficient atomic CPU instructions in the implementation. Consequently, the proposed method scales linearly with the number of *k*-mers and genomes, something that would not be possible if *k*-mer frequencies were considered. In future work, we will explore ways to incorporate *k*-mer frequencies, while preserving the scalability of our method. This new type of model will allow the detection of *k*-mers at multiple genomic loci, which could prove important for modeling phenotypes that are affected by structural variations, such as copy number variations.

## Abbreviations

PCR: Polymerase chain reaction
SCM: Set Covering Machine
SNP: Single nucleotide polymorphism
L1SVM: L_1_-norm regularized Support Vector Machine
L2SVM: L2-norm regularized Support Vector Machine
LinSVM: Linear kernel Support Vector Machine
MEGA: Macrolide efflux genetic assembly
PolySVM: Polynomial kernel Support Vector Machine
RRDR: Rifampicin resistance-determining region

## Availability of supporting data

The data sets supporting the results of this article are available in the GenBank and EMBL repositories: PRJEB2632, PRJEB11776, PRJEB7281, PRJNA264310.

Kover, the out-of-core implementation of our method, is open-source and available at http://github.com/aldro61/kover.

## Competing interests

The authors declare that they have no competing interests.

### Author contributions

AD, FL, MM and SG designed the algorithmic extensions to the Set Covering Machine algorithm. AD, FL and MM derived the sample compression bound for the Set Covering Machine. AD designed the out-of-core implementation of the Set Covering Machine. AD, FL, JC, MM and SG designed the experimental protocols and AD conducted the experimentations. AD, JC, MD and SG evaluated the biological relevance of the models. MD acquired the data and prepared it for analysis. AMB and VL acquired and provided the C. difficile genomes and the associated antibiotic resistance data. AD, FL, JC, MD, MM, MT and SG wrote the manuscript. All authors have read and approved the final manuscript.

### Additional data files

The following additional data are available with the online version of this paper. Additional data file 1 is a table containing the antibiotic resistance classifications for the *Clostridium difficile* genomes of Dr. Loo and Dr. Bourgault [EMBL:PRJEB11776].

## Acknowledgements

The authors acknowledge Dr. Eric Audemard, Dr. Sebastien Boisvert, Maia Kaplan, Dr. Sylvain Moineau, Dr. Jean-Louis Plouhinec and Dr. Paul H. Roy for helpful comments and suggestions. Computations were performed on the Colosse supercomputer at Université Laval (resource allocation project: nne-790-ae), under the auspices of Calcul Québec and Compute Canada. AD is recipient of an Alexander Graham Bell Canada Graduate Scholarship Doctoral Award of the National Sciences and Engineering Research Council of Canada (NSERC). This work was supported in part by the NSERC Discovery Grants (FL; 262067, MM; 122405) and an award to MT from the Ministére de l’enseignement supérieur, de la recherche, de la science et de la technologie du Québec through Génome Québec. JC acknowledges the Canada Research Chair in Medical Genomics. AMB and VL acknowledge the Consortium de Recherche sur le Clostridium difficile, which consists of the following partners: Fonds de la Recherche en Santé du Québec, Canadian Institutes of Health Research, Ministre de la Santé et des Services Sociaux du Québec, Institut National de Santé Publique du Québec, Health Canada, Centre Hospitalier de l’Université de Montréal, McGill University Health Centre, CHU de Québec, and CHU de Sherbrooke.

## Additional files

### Additional file 1 — Appendix

The appendix contains supporting information and detailed results that are not essential to the reader’s understanding of the key findings of this study.

### Additional file 2 — Supplemental Tables

Table S1: A detailed overview of the sequencing data used in this study; Table S2: A detailed overview of the antibiotic resistance datasets used in this study; Table S3: An analysis of overfitting for the Set Covering Machine and support vector machines with linear and polynomial kernels; Table S4: Average and standard deviation of the sensitivities measured on the testing set for 10 random partitions of each dataset; Table S5: Average and standard deviation of the specificities measured on the testing set for 10 random partitions of each dataset; Table S6: Average and standard deviation of the error rates measured on the testing set for 10 random partitions of each dataset.

### Additional file 3 — Supplemental Figures

Figure S1: An illustration of the resistance models generated for each antibiotic.

